# The usefulness of sparse k-means in metabolomics data: An example from breast cancer data

**DOI:** 10.1101/2022.02.05.479235

**Authors:** Misa Goudo, Masahiro Sugimoto, Satoru Hiwa, Tomoyuki Hiroyasu

## Abstract

In processing metabolomics data, multidimensional quantitative data from thousands of metabolites are often sparse, that is, only a small fraction of metabolites are relevant to the phenotype of interest. Clustering is therefore used to discover subtypes from omics data. Sparse processing, which selects important metabolites from the total omics data, is an effective clustering technique. This study investigated the effectiveness of sparse k-means for metabolomics data. Specifically, sparse k-means was used to cluster blood lipid metabolite data of breast cancer patients in two studies: (1) before and after menopause, and (2) pre- and postoperative chemotherapy. In both cases, sparse k-means showed comparable discrimination accuracy with fewer metabolites than k-means. Furthermore, when the L1 norm values were varied, no significant changes were observed. The mean silhouette coefficients of sparse k-means and k-means were (1) 0.38 ± 0.14 (S.D.) and 0.17 ± 0.01, (2) 0.38 ± 0.07 and 0.17 ± 0.01, indicating that feature selection using sparse k-means can improve clustering results. In addition, metabolite selection using sparse k-means was consistent regardless of the test data or the constrained value of the L1 norm, indicating robustness.

## 1 Introduction

Recent advances in omics technologies have made it possible to comprehensively quantify molecules in living organisms and generate extensive data, such as proteomes and transcriptomes [1]. By observing changes in hundreds to thousands of molecular-level networks, it has become possible to comprehensively understand the etiology and pathogenesis of various diseases [2], search for diagnostic and predictive markers, and search for therapeutic targets based on patterns of multiple molecules rather than single molecules [3]. Clustering is used to interpret multidimensional quantitative omics data, where highly similar data can be extracted as clusters. It has been used to search for new subtypes of diseases and classify data using molecular aggregates [4]. Clustering is often used to identify subtypes of a particular disease by integrating methylation and genomic information with the transcriptome [5, 6]. Whether in single-omics or multi-omics, clustering is considered one of the most effective methods to consolidate various information inherent in the data.

There are various clustering methods, of which the most widely used are hierarchical clustering and k-means clustering, unsupervised learning methods [7, 8]. According to a predefined distance measure, k-means clustering classifies practical information into k clusters [8]. A similar technique is Partitioning Around Medoids (PAM), which introduces dissimilarity (dissimilarity matrix). Other single-layer neural network methods such as Self-Organizing Maps (SOM) [9] have also been developed. These methods classify the similarity of samples and rank the importance of observed variables based on their clustering contribution and use the top-ranked variables for biochemical considerations and markers. However, all of these methods use all observed variables; thus, many variables are involved in extracting information for each cluster, which presents computational challenges. In contrast, sparse k-means, an improved version of k-means [10], allows clustering with fewer observables, eliminating information that has little relevance to the cluster, leaving only important variables for cluster formation.

Metabolomics is a comprehensive and simultaneous molecular profiling method that captures metabolic data. Nuclear magnetic resonance (NMR) and mass spectrometry (MS) are widely used in metabolomics. The instruments used in these techniques are becoming increasingly sensitive, enabling the simultaneous measurement of hundreds to thousands of metabolites. The high sensitive mass spectrometers have enabled the quantification of a wide variety of metabolites, which increased the dimension of the observed information [11, 12]. Multivariate analysis, such as principal component analysis (PCA), is a standard method to obtain a complete picture of measured metabolites [13]. Predictive models, such as partial least squares regression (PLS-DA) and two-way orthogonal partial least squares (O2PLS), are widely used to rank molecules that contribute to classification and are correlated with quantitative values [14]. K-means metabolic data analysis has been applied in various fields. For example, in an Irish cohort, triglycerides and glucose in fasting serum samples were clustered using k-means to identify individual dietary habits [15]. In another study, k-means clustered metabolic profiles from liquid chromatography-tandem mass spectrometry (LC-MS/MS) data were used from red wine [16].

Thus, to discover new biochemical mechanisms and establish molecular markers, it is necessary to extract representative metabolites among the metabolites detected. Furthermore, if the discrimination accuracy is similar or improved using only a subset of features compared to all features, these subset metabolites can explain the focused phenotype. Therefore it is essential to extract a subset of metabolites for facilitating the discovery of new biochemical mechanisms and molecular markers.

Among the clustering methods, sparse k-means can identify essential features simultaneously with clustering. Witten and Tibshirani developed an alternative method that identifies features using lasso-type penalties based on conventional k-means [10]. Sparse k-means can also be effective for metabolomics data; however, there are few metabolomics studies that utilize sparse k-means [17]. Therefore, it is necessary to investigate the impact of each clustering process on the accuracy of the data and establish an analytical protocol to evaluate the applicability of sparse k-means in metabolomic data analysis.

This study conducted two comparisons of sparse k-means and k-means using quantitative blood lipid data from breast cancer patients: one for premenopausal and menopausal, and another for preoperative and postoperative chemotherapy.

## 2 Materials and Methods

### 2.1 Mini-review of existing methods

Two unsupervised clustering methods, k-means and sparse k-means, were compared. The k-means clustering method is a well-known algorithm [18]. The number of clusters (*k*), which is the number of clusters to be extracted, was pre-specified by a data analyst. The clusters were determined by minimizing the within-cluster sum of squares (WCSS). This operation is synonymous with maximizing the between-cluster sum of squares (BCSS). In contrast, sparse k-means is an algorithm for clustering using features selected during adaptation [10], where a weight of *≥*0 is assigned to each metabolite. The sparse k-means is a function of the sum of the squares between the clusters multiplied by the weights of the variables, as shown in Equation 1. The clusters are determined by maximizing this sum [10].

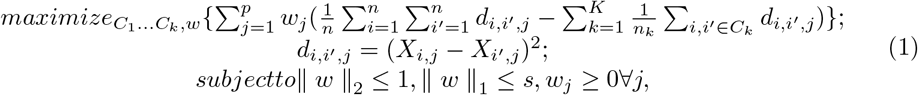

where *X* is an *n* (sample) *×p* (variable) matrix, *i* is the number of samples, *j* is the number of variables, *w* is the weight of the variable, *w*_*j*_ is the weight of the *j*th variable, *k* is the number of clusters, *n*_*k*_ is the number of samples in the *k*th cluster number, and *C*_*k*_ is the *k*th cluster. The parameter *s* is the sum of the weights of the features. If *s* is large, more features are selected, and if *s* is small, fewer features are selected.

### 2.2 Evaluation indicators and data used in this study

#### 2.2.1 Evaluation indicators

Four evaluation metrics were calculated: accuracy, sensitivity, and specificity to validate the data classification and the silhouette coefficient to measure the cohesiveness of the clusters. The first three metrics were calculated from the mixing matrix. Accuracy was calculated by(true positive [TP] + true negative [TN]) / (TP + TN + false positive [FP] + false negative [FN]), sensitivity by TP / (TP + TN), and specificity by TN / (TN + FP). The silhouette coefficient is an index for evaluating clustering performance, based on the idea that clustering results are condensed within clusters and distant in different clusters. The method used to calculate the silhouette coefficient is given by Equation 2.

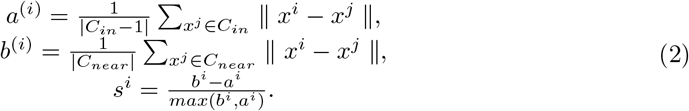

Here, *a*^(*i*)^ is the average distance from sample *i* to all samples in the same cluster (*C*_*in*_) and *b*^(*i*)^ is the average distance from sample *i* to all samples of the closest cluster (*C*_*in*_). The silhouette coefficient is calculated from −1 to 1, where 1 indicates that the sample is farther away from neighboring clusters and −1 indicates that the sample is far from the center of its cluster, which means that the sample may be in the wrong cluster. A silhouette coefficient of 0 indicates that the sample is on or near the decision boundary. The average silhouette coefficients for all data were used to assess the validity of the clustering [19].

#### 2.2.2 Data selection

The MTBLS92 dataset from MetaboLights (https://www.ebi.ac.uk/metabolights/)was used in this study. This dataset was obtained from a multicenter, randomized, phase III trial in which breast cancer patients were randomly assigned to one of the following preoperative neoadjuvant chemotherapies (NACs).

1. 4*×*epirubicine+cyclophosphamide(EC)*→*docetaxel(D)
2. 4*×*EC*→*4*×*D/capecitabine(C)
3. 4*×*EC*→*4*×*D*→*C

Blood samples were collected before NAC (baseline, BL) and in the non-fasting state after NAC at the time of surgery. Serum lipid levels were quantified by liquid chromatography-mass spectrometry (LC) and fatty acids by gas chromatography (GC), and a flame ionization detector (FID) detector. Substances were quantified based on the peak intensity ratio and using internal standards for each pretreatment and were identified by a library search developed internally. The dataset consisted of 253 individuals and 240 metabolites. We refer the reader to the original paper for details of the study design, sample collection, and processing [20]. The number of samples used in this study is listed in Table 1.

**Table 1.**
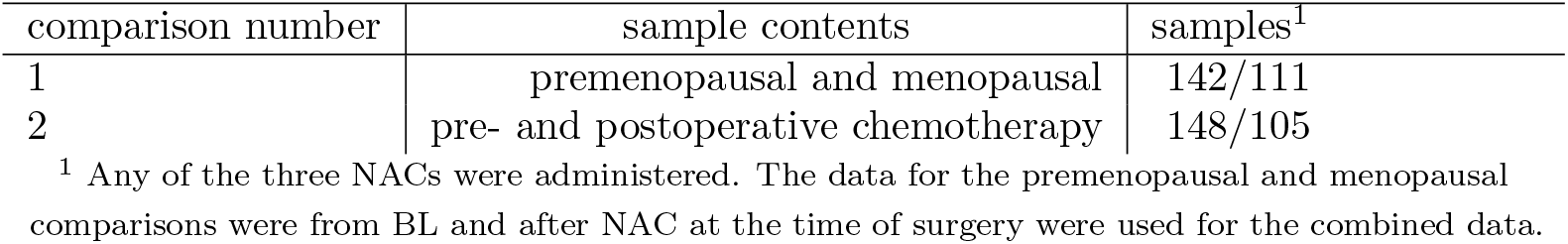
Number of samples

### 2.3 Clustering process

Hilvo et al. showed a difference in the metabolite profiles of patients receiving NAC before and after treatment. They also showed a difference in the menopausal period [20]. This study used two-class clustering to separate pre- and postoperative chemotherapy and premenopausal and menopausal patients based on these results. The data was divided into two datasets: one for class determination and the other for prediction to perform two-class clustering. K-means and sparse k-means clustering were applied to the datasets used for class determination, and the centers of the two clusters were determined. The sum of the weights of the features (*s*), the option with the highest accuracy when combined with the teacher data, was adopted. The most represented label determined the label of each cluster in each respective cluster. The labels of close centers categorized the dataset for prediction. Finally, the accuracy, sensitivity, specificity, and average of the silhouette coefficients were calculated for performance evaluation. In the premenopausal and menopausal classification, values were assigned as positive and negative, respectively; in the pre- and postoperative chemotherapy clustering, values were assigned as positive and negative, respectively. All statistical analyses used in this study were based on the Welch’s t-test (*P <* 0.05). The dataset was split into two datasets by a stratified extraction method to determine 4/5 and predict 1/5 classes. The dataset was split ten times, and the same dataset was used for each method to minimize bias. Missing values in the data were interpolated using half of the mean value of each variable. In addition, the values were log2-transformed and normalized per sample, and then the normalized values per variable were used. The default number of initial searches for each method was 10, and the number of clustering attempts was set to 100. Sparse k-means requires the sum of the weights of the features (denoted by *s* in Equation 1). In this study, we did not determine it uniquely but varied it in increments of 0.48 to divide the data into 40 parts between 1.1 and 20. The minimum value of *s* is 1.1, and at *s* = 20, the results were stable in preliminary trials. Out of ten trials, the variables whose features were nonzero and were ranked according to the most significant coefficient, and the total value of the ten rankings was calculated. For example, ten was calculated if the variable was ranked first ten times.

### 2.4 Software implementation

K-means and sparse k-means were computed using R (v 4.0.3; R Development Core Team, Vienna, Austria). K-means was computed using the standard R package “stats” (v 4.0.3). Sparse k-means was computed using the R package “sparcl” (v 1.0.4) [21]. The program will be released upon request.

## 3 Results

### 3.1 Differences in the selected features depend on the constraints of the weights in sparse k-means

Once the features are selected, the data can be represented in a lower dimension, which is necessary for determining metabolites for practical applications. The sparse k-means algorithm is characterized by features becoming sparse after clustering. However, obtaining a sparse result depends on the hyperparameter setting of the sparse k-means, that is, the sum of the weights of each feature (*s*). By changing *s*, the weights attached to each feature will change, also changing the number of features with zero weight. Therefore, in sparse k-means, the feature selected varies greatly depending on *s*.

Figure 1 shows the relationship between *s* and the number of metabolites selected using sparse k-means. For high values of *s*, the selected metabolites included all metabolites (240), indicating that selected features were not sparse. In the two-class clustering dividing premenopausal and menopausal classes, all metabolites were selected when *s ≥* 12.73, indicating that sparsity was not reached. When *s <* 12.73 was sparse, i.e. only some metabolites were selected. Two-class clustering based on pre- or postoperative chemotherapy also yielded the same results as those based on menopausal status. In both clustering methods, features were limited with *s <* 12.73, confirming the sparsity of selected features.

**Figure 1.**
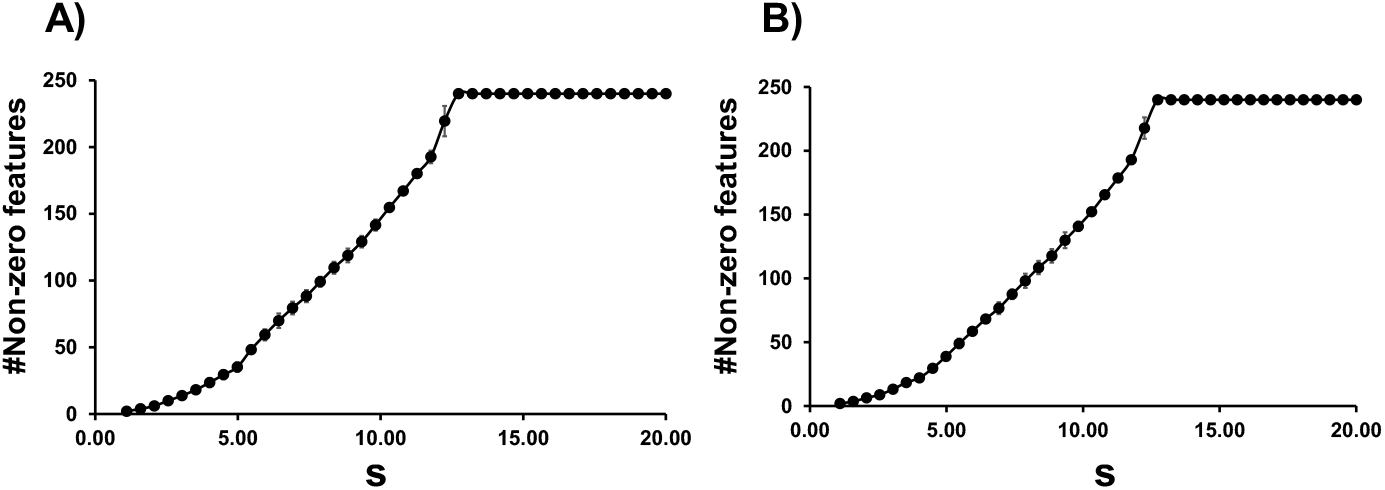
Relationship between the sum of the weights of the features (*s*) and the number of variables with nonzero weights for two-class clustering. Results for (**A**) premenopausal and menopausal, and (**B**) pre- and postoperative chemotherapy. The x-axis shows *s*, and the y-axis shows the number of variables with nonzero weights. Error bars indicate standard deviations.

### 3.2 Comparison of sparse k-means and k-means classification accuracy

This study used accuracy, sensitivity, and specificity as indices for class classification. In addition, average silhouette coefficients were used as indicators of cluster cohesion. The classification accuracy was determined based on the distance between the centers of the clusters. Figure 2 shows the evaluation indices for the sparse k-means and k-means classification, resulting in a higher accuracy than the clusters’ data. In the two-class clustering separating premenopausal and menopausal classes, the mean silhouette coefficients of sparse k-means and k-means were 0.38 ± 0.14 (S.D.) and 0.17 ± 0.01, respectively, indicating a significant difference. No significant differences were found for the other indices. Similarly, in the two-class clustering separating pre- and postoperative chemotherapy classes, the mean silhouette coefficients of sparse k-means and k-means were 0.38 ± 0.07 and 0.17 ± 0.01, respectively, indicating a significant difference. Otherwise, no significant differences were observed. Figure 3 shows the accuracy of the created clusters compared with the learning data. The labels of the learning data were assigned to the center of the clusters. There was a significant difference between the two classifications separating premenopausal and menopausal classes, with a sparse k-means of 0.62 ± 0.01 and k-means of 0.60 ± 0.01. In the two-class clustering separating the pre- and postoperative chemotherapy classes, the sparse k-means and k-means were 0.57 ± 0.02 and 0.56 ± 0.02, respectively, and no significant difference was obtained. A variable (row) × sample (column) matrix was created for each TP, FP, FN, and TN result obtained from the mixing matrix. Figure 4 shows the matrices for k-means and sparse k-means clustering. For each comparison, the sparse k-means selected metabolites that characterized the cluster better than k-means because the heat map colors differed between the labels. This suggests that metabolites selected by sparse k-means capture the characteristics of the clusters.

**Figure 2.**
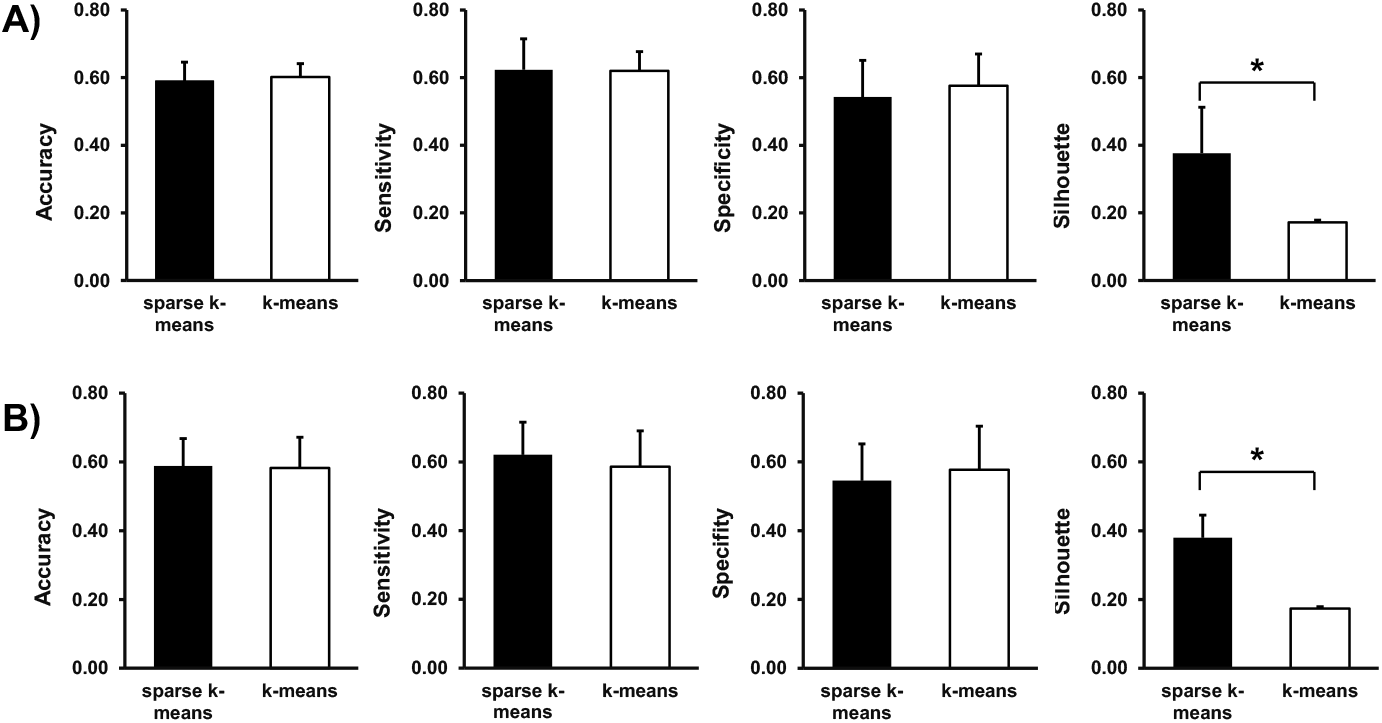
Comparison of evaluation indices for two-class clustering. Results for (**A**) premenopausal and menopausal, and (**B**) pre- and postoperative chemotherapy. From left to right, both panels show the accuracy, sensitivity, specificity, and silhouette coefficient. Error bars indicate standard deviation obtained from 10 trials. *\*P <* 0.05 (Welch’s t-test).

**Figure 3.**
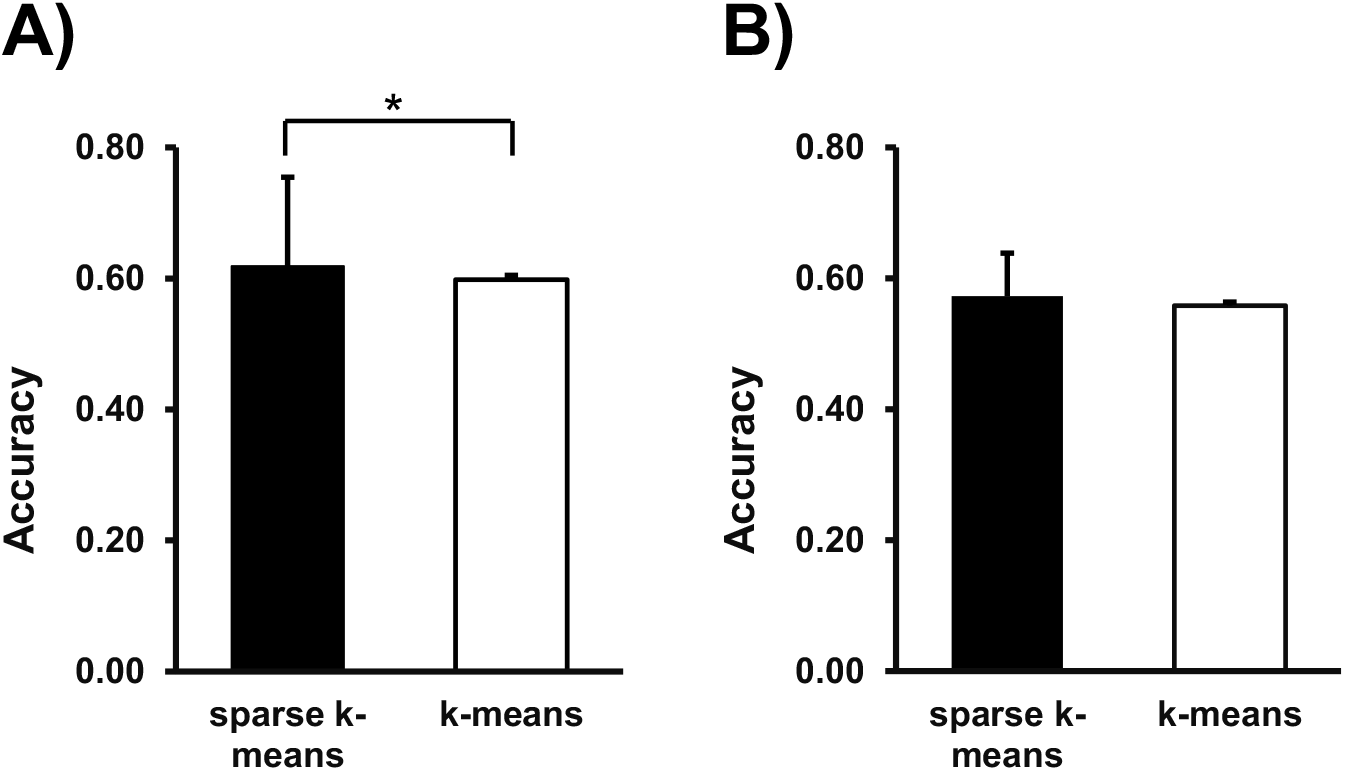
Accuracy of cluster labeling for (**A**) premenopausal and menopausal, and (**B**) pre- and postoperative chemotherapy two-class clustering. Error bars indicate standard deviation obtained from 10 trials. *\*P <* 0.05 (Welch’s t-test).

**Figure 4.**
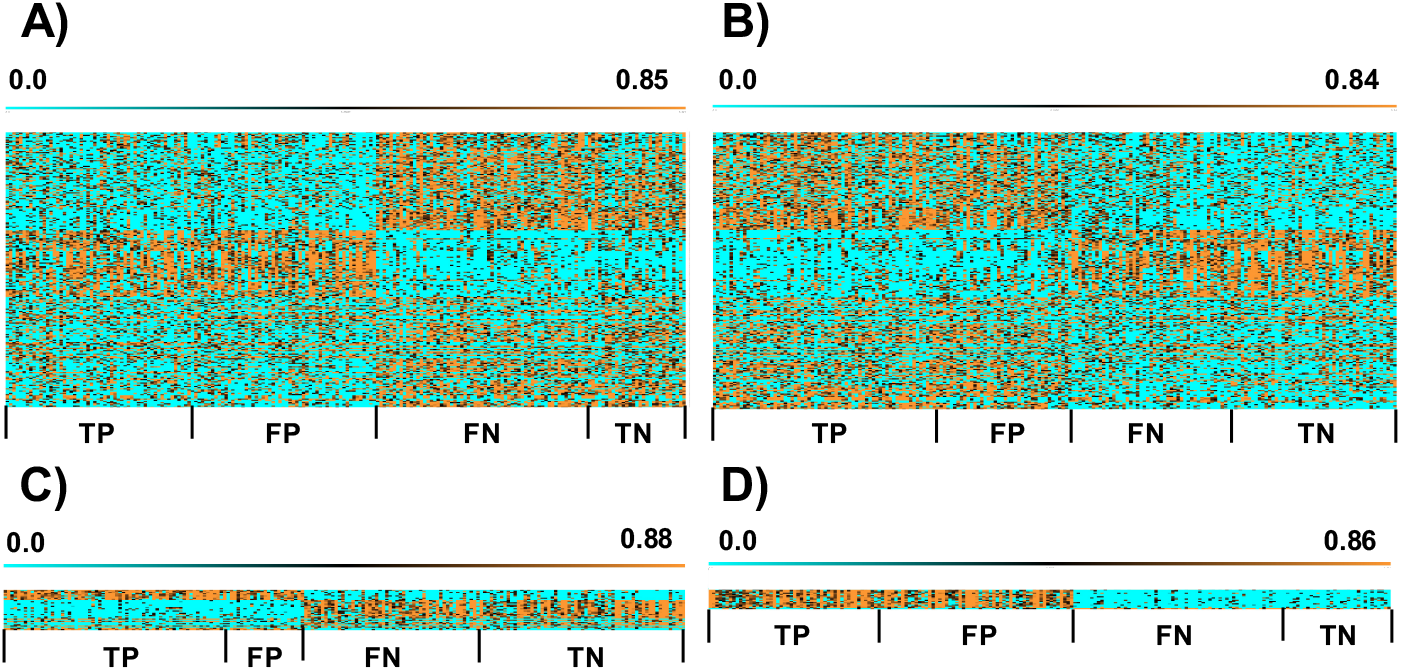
Matrices for (**A, C**) premenopausal and menopausal, and (**B, D**) pre- and postoperative chemotherapy two-class clustering. In (**A**) and (**B**), k-means produced a matrix of 240 (metabolites) *×* 202 (samples); in (**C**), sparse k-means (*s* = 4.49) produced a matrix of 35 (metabolites) *×* 202 (samples); in (**D**), sparse k-means (*s* = 3.52) produced a matrix of 18 (metabolites) *×* 202 (samples).All panels show, from left to right, True positive (TP), False Positive (FP), False Negative (FN), and True Negative (TN), where premenopausal and preoperative chemotherapy are considered positive and menopausal and postoperative are considered negative. The scale value indicates the value of the metabolic profiles.

### 3.3 selected metabolites

Table 2 and 3 show the list of metabolites selected by sparse k-means for comparison 1 (premenopausal and menopausal) and comparison 2 (pre- and postoperative chemotherapy), respectively. In Table 2, unidentified 42 (unidentified substance) was selected nine times out of 10, triglycerides (TG; 14:0/16:0/18:1) were selected seven times, and the total ranking value was the smallest among all metabolite profiles. In Table 3, the median number of variables was 33.5, with a minimum of 6 and a maximum of 89 in 10 trials; TG (14:0/16:0/18:0), TG (16:0/16:0/18:0), phosphatidylcholine (PC), and sphingomyelin (SM) were included in the majority of trials. All other molecules were selected at least twice, with the exception of ceramide, (Cer; d18:1/17:0), TG (16:0/18:1/20:1), TG (18:0/18:1/18:1), PC (34:1e) | PE (phosphatidylethanolamine (37:1e), and a few unidentified substances.

**Table 2.** Variables selected in comparison 1 (top 15)

| metabolite | number of times selected | total ranking value |
| --- | --- | --- |
| unidentified_42 | 9 | 46 |
| TG(14:0/16:0/18:1) | 7 | 24 |
| TG(48:2) | 6 | 175 |
| TG(16:0/18:1/16:0) | 6 | 146 |
| TG(14:0/16:0/18:0) | 6 | 134 |
| TG(16:0/16:0/18:0) | 6 | 159 |
| TG(46:1) | 6 | 192 |
| unidentified_18 | 6 | 258 |
| TG(14:0/16:0/16:0) \| TG(16:0/18:0/12:0) | 6 | 267 |
| SM(d18:1/22:1) | 6 | 281 |
| unidentified_47 | 6 | 278 |
| TG(14:0/18:1/18:1) \| TG(16:0/16:1/18:1) | 6 | 273 |
| unidentified_53 | 6 | 456 |
| TG(49:1) | 5 | 190 |
| TG(16:0/18:0/18:1) | 5 | 188 |

**Table 3.** Variables selected in comparison 2 (top 15)

| metabolite | number of times selected | total ranking value |
| --- | --- | --- |
| TG(14:0/16:0/18:1) | 10 | 28 |
| TG(14:0/16:0/18:0) | 10 | 38 |
| unidentified_42 | 10 | 41 |
| TG(16:0/16:0/18:0) | 10 | 67 |
| unidentified_47 | 10 | 91 |
| TG(14:0/16:0/16:0) \| TG(16:0/18:0/12:0) | 9 | 66 |
| TG(46:1) | 8 | 63 |
| TG(16:0/18:1/16:0) | 8 | 90 |
| TG(48:2) | 8 | 108 |
| TG(16:0/18:0/18:1) | 8 | 138 |
| unidentified_18 | 8 | 176 |
| TG(47:1) | 8 | 179 |
| Custom adduct of 796,742 m/z - TG(14:0/16:0/16:0)+TG(16:0/18:0/12:0) | 7 | 111 |
| unidentified_71 | 7 | 125 |
| TG(44:1) | 7 | 146 |

## 4 Discussion

### 4.1 Clustering Accuracy

This study used sparse k-means and k-means to investigate classification accuracy and cluster cohesion by performing two-class clustering using two comparisons, one for premenopausal and menopausal and the other for pre- and postoperative chemotherapy. There was no significant difference in classification accuracy between sparse k-means and k-means, but sparse k-means showed a significant difference in cluster cohesion due to variable selection. Sparse k-means was more in line with the teacher data in the premenopausal and menopausal comparison than k-means. This result suggests that sparse k-means can perform clustering and feature selection, providing more information than k-means. Originally, it was necessary to measure the gap statistics to determine *s* [10, 17, 22]. In this study, this was not considered. In addition, we did not compare sparse k-means with other methods, such as PLS-DA. Future studies may investigate the effects of different clustering methods in metabolite selection. Gal et al. compared the metabolic signatures of breast cancer obtained using five unsupervised learning methods, including sparse k-means [17]. They determined the s in sparse k-means using the gap statistic, whereas, in this paper, we compared the differences in metabolites with changes in *s*. We also examined the accuracy of the classification based on its proximity to the center of the cluster. The k-means method uses all the features for clustering, so all the observed metabolites can be divided into similar groups. On the other hand, the sparse k-means method is feature-selective to extract specific metabolites within similar groups.

### 4.2 Metabolites selected by sparse k-means

Although the number of variables selected by sparse k-means varied among trials, most variables were selected in multiple trials. In particular, only triglycerides were selected in the smallest number of trials in the pre- and postoperative comparisons. In breast cancer patients treated with NAC, the blood lipid profile remains unchanged, but changes in thiobarbituric acid reactive substances (TBARS) and superoxide dismutase (SOD) indicate that oxidative stress in the body is significantly altered [23], resulting in a change in TG levels. Epirubicin, an anthracycline anticancer drug, was included in the NAC. It has been reported that some SMs may contribute to the activation efficiency of anthracycline uptake [24]. PC has been shown to protect against peripheral neurotoxicity, one of the side effects caused by docetaxel [25]. Changes in the blood profile with NAC treatment are also speculated. In breast cancer subtypes, triple-negative tumors contain more specific PCs [26], suggesting that the baseline may differ in each subtype rather than uniformly changing in all cases. In contrast, there are reports that blood PC in breast cancer patients decreases during anticancer drug treatment [27]. PC itself can be used as a marker for predicting cervical cancer susceptibility to NAC [28]. Therefore, changes in the blood profile may vary from individual to individual.

## 5 Conclusions

In metabolomics data analysis, k-means and hierarchical clustering are used for discriminating multiple groups. However, it is difficult to examine metabolic characteristics in a vast number of metabolites. Therefore, clustering with a few metabolites and feature selection is essential for analysis. Although sparse k-means, which simultaneously performs feature selection and clustering, is one method to solve this problem, it has not been widely adopted for metabolomics analysis. This study investigates the effect of constraint values on the weights of sparse k-means when applied to metabolomics data and examines its usefulness for future analysis. In this study, two breast cancer metabolomics data classifications were conducted using k-means and sparse k-means. No statistically significant difference in classification accuracy was found between sparse k-means and k-means clustering. The sparse k-means method selected variables that explained the clusters by setting appropriate weight constraint values. In addition, the selected metabolites were suitable for discriminating the given problems. Because feature selection can be made simultaneously with clustering, the sparse k-means method is useful for analyzing metabolomics data.

